# Structures of the *Pseudomonas aeruginosa* MlaC–MlaD complexes reveal a conformational switch mediated by the C-terminal helix of MlaC

**DOI:** 10.64898/2026.05.15.725309

**Authors:** Daiki Matsumoto, Sayaka Ozu, Yasunori Watanabe

## Abstract

Gram-negative bacteria maintain an asymmetric outer membrane that protects cells from environmental stresses and antibiotics. The maintenance of lipid asymmetry (Mla) pathway contributes to outer membrane lipid homeostasis through phospholipid transport between the outer and inner membranes. Although the periplasmic lipid carrier MlaC is thought to transfer phospholipids to the inner membrane MlaFEDB transporter via the hexameric protein MlaD, the molecular mechanism underlying this process remains unclear. Here we show crystal structures of two distinct MlaC–MlaD complexes from *Pseudomonas* aeruginosa that reveal distinct conformational states of MlaC. In these structures, an ordered conformation of the C-terminal α8 helix of MlaC positions MlaC distally from the central pore of the MlaD hexamer and limits accessibility of the lipid-binding cavity, whereas partial disordering of the α8 helix allows closer association with the MlaD hexamer and increased exposure of the cavity. Structure-based biochemical analyses further demonstrate that the C-terminal region negatively regulates MlaC– MlaD interaction while stabilizing phospholipid binding. These findings identify the C-terminal α8 helix as a conformational switch that couples MlaC positioning with lipid cavity accessibility, providing structural insight into phospholipid transfer at the MlaC–MlaD interface.

## Introduction

Gram-negative bacteria are surrounded by two membranes, the outer and inner membranes, forming a protective envelope important for viability and environmental adaptation. The outer membrane (OM) is an asymmetric lipid bilayer composed of phospholipids in the inner leaflet and lipopolysaccharides (LPS) in the outer leaflet^1^. This asymmetric organization is critical for maintaining the permeability barrier function of the OM, thereby protecting bacteria from toxic compounds such as antibiotics and detergents^2^.

One of the major pathways implicated in maintaining OM lipid asymmetry is the maintenance of lipid asymmetry (Mla) pathway, a widely conserved pathway in Gram-negative bacteria that is important for OM lipid asymmetry and membrane integrity^3-5^. The Mla system is a multi-component machinery composed of the OM protein MlaA in complex with an osmoporin OmpF/C^6,7^, the periplasmic lipid binding protein MlaC^8^, and the inner membrane (IM) ABC transporter complex MlaFEDB^9^. The MlaFEDB complex is thought to couple ATP hydrolysis to phospholipid transfer. Together, these components are thought to mediate the transport of phospholipids between the OM and IM, thereby contributing to lipid homeostasis. Although the directionality of transport has been debated^10-12^, accumulating evidence suggests that the Mla pathway plays a key role in retrograde phospholipid transport from the OM to the IM^13-15^. In *Escherichia coli*, disruption of the Mla pathway leads to defects in OM lipid asymmetry and increased sensitivity to detergents and antibiotics, consistent with impaired OM integrity^3,13^. In several Gram-negative pathogens, Mla pathway defects have also been linked to reduced virulence or defects in virulence-associated phenotypes, including *Pseudomonas aeruginosa*^16-21^.

MlaC interacts with both the OM-associated MlaA–OmpF/C complex and the IM-associated MlaFEDB complex, suggesting that it functions as a soluble periplasmic shuttle for phospholipids^8^. Previous structural studies have revealed that MlaC adopts a mixed α/β fold containing a deep hydrophobic cavity capable of accommodating phospholipid molecules^8^. Crystal structures of MlaC homologs from several bacterial species have further revealed open and closed conformations of the lipid-binding cavity, suggesting a dynamic mechanism for lipid loading and release^22,23^. In addition, structural studies of the MlaFEDB complex from multiple bacterial species have provided insights into how phospholipids are inserted into the inner membrane^13,24-28^. More recently, cryo-electron microscopy (cryo-EM) analysis of the *E. coli* MlaFEDB–MlaC complex together with AlphaFold2-predicted structural models suggested that MlaC can adopt multiple docking modes on the MlaD hexamer^29^. In addition, cryo-EM structures of the *E. coli* MlaC–MlaD complex provided structural insights into how MlaC associates with the MlaD hexamer^30^. Despite these advances, the structural basis underlying phospholipid transfer at the MlaC–MlaD interface remains incompletely understood.

In this study, we determined the crystal structures of the MlaC–MlaD complexes from *P. aeruginosa* and investigated the role of the C-terminal α8 helix of MlaC in lipid binding and interaction with MlaD. Our structural analysis reveals distinct conformational states of MlaC in complex with the MlaD hexamer. We further show that conformational rearrangement of the C-terminal α8 helix modulates both the orientation of MlaC and the accessibility of its lipid-binding cavity. Biochemical analyses demonstrate that the C-terminal region negatively regulates MlaC– MlaD interaction while stabilizing phospholipid binding. Together, our findings provide new insights into the molecular mechanism of phospholipid transfer in the Mla pathway and support a model in which the C-terminal region of MlaC regulates phospholipid transfer.

## Results

### Crystal structure of the MlaC–MlaD^ΔN36^ complex from *P. aeruginosa*

To gain structural insight into the phospholipid transfer between MlaC and MlaD, we sought to determine the structure of the MlaC–MlaD complex from *P. aeruginosa*. The mature periplasmic form of MlaC (residues 23–215), lacking the N-terminal signal peptide, was used for crystallization and is hereafter referred to as PaMlaC. Two MlaD variants (residues 31–157 and 37–157), lacking the N-terminal signal peptide and the transmembrane helix, were constructed and are hereafter referred to as PaMlaD^ΔN30^ and PaMlaD^ΔN36^. An in vitro pull-down assay showed that both PaMlaD^ΔN30^ and PaMlaD^ΔN36^ interacted with PaMlaC (Supplementary Fig. 1). We therefore used both PaMlaD constructs for crystallization with PaMlaC and obtained well-diffracting crystals of the PaMlaC–PaMlaD^ΔN30^ and PaMlaC–PaMlaD^ΔN36^ complexes. We then determined the crystal structure of the PaMlaC–PaMlaD^ΔN36^ complex by the molecular replacement method using the structures of *P. aeruginosa* MlaC (PDB code: 6HSY)^22^ and the *P. aeruginosa* MlaFEDB complex (PDB code: 7CH9)^28^ as the search models. The structure was refined to 2.56 Å resolution (Table 1). Consistent with previous studies^8,28^, six PaMlaD^ΔN36^ molecules (PaMlaD_1–6_) assembled into a ring-shaped hexamer, with their C-terminal helices (α1_1_–α1_6_) forming a central pore (Fig. 1A). In the asymmetric unit, three PaMlaC molecules (PaMlaC_1–3_) were bound to the PaMlaD^ΔN36^ hexamer in a threefold symmetric arrangement (Fig. 1A). Specifically, PaMlaC_1_, PaMlaC_2_, and PaMlaC_3_ interacted with PaMlaD_1_, PaMlaD_3_, and PaMlaD_5_, respectively, while also contacting the C-terminal helices of PaMlaD_2_, PaMlaD_4_, and PaMlaD_6_ (Fig. 1B). Structural comparison of PaMlaC_1–3_ with the previously reported structure of *P. aeruginosa* MlaC (PDB code: 6HSY) revealed that the overall structure was largely unchanged in this complex, with r.m.s.d. values of 0.62–0.75 Å over 167–184 Cα atoms. PaMlaC_1_ and PaMlaC_2_ each contained electron densities consistent with two phospholipid molecules in their cavities (Fig. 1C and Supplementary Fig. 2), whereas no interpretable electron density was observed in the cavity of PaMlaC_3_. Thin-layer chromatography (TLC) analysis showed that purified PaMlaC contained both phosphatidylethanolamine (PE) and phosphatidylglycerol (PG) derived from *E. coli* membranes (Fig. 1D). Because the electron density for the head group was poorly defined, the phospholipid species could not be unambiguously assigned as PE or PG. Therefore, only the phospholipid moiety excluding the head group was modeled. The phosphate groups of the phospholipids in PaMlaC were oriented toward the central pore of the PaMlaD hexamer. This arrangement is consistent with a model in which PaMlaC delivers phospholipids to the central pore of the PaMlaD hexamer.

**Table 1.** Data collection and refinement statistics.

|  | MlaC–MlaD<sup>ΔN36</sup><br>[PDB ID 26TP] | MlaC–MlaD<sup>ΔN30</sup><br>[PDB ID 26TQ] |
| --- | --- | --- |
| **Data collection** |  |  |
| Space group | *P*1 | *P*4<sub>1</sub>2<sub>1</sub>2 |
| Cell dimensions |  |  |
| *a*, *b*, *c* (Å) | 61.71, 88.64, 88.81 | 125.16, 125.16, 178.90 |
| $\alpha$, $\beta$, $\gamma$ (°) | 64.65, 72.37, 72.28 | 90.00, 90.00, 90.00 |
| Resolution (Å) | 50.0–2.56 (2.71–2.56) | 50.0–3.46 (3.67–3.46) |
| *R*<sub>sym</sub> | 0.317 (1.178) | 0.410 (2.966) |
| *R*<sub>meas</sub> | 0.393 (1.475) | 0.422 (3.089) |
| *I* / $\sigma I$ | 3.1 (0.8) | 7.7 (0.9) |
| Completeness (%) | 97.0 (95.5) | 99.8 (99.0) |
| Redundancy | 2.6 (2.6) | 17.5 (12.9) |
| CC<sub>1/2</sub> | 0.878 (0.358) | 0.998 (0.476) |
| **Refinement** |  |  |
| Resolution (Å) | 47.4–2.56 | 49.5–3.46 |
| No. reflections | 50503 | 17545 |
| *R*<sub>work</sub> / *R*<sub>free</sub> | 0.249 / 0.303 | 0.300 / 0.357 |
| No. atoms |  |  |
| Protein | 8964 | 8451 |
| Ligand/ion | 176 | - |
| Water | 71 | - |
| *B*-factors |  |  |
| Protein | 66.34 | 102.22 |
| Ligand/ion | 85.24 | - |
| Water | 54.12 | - |
| R.m.s. deviations |  |  |
| Bond lengths (Å) | 0.003 | 0.003 |
| Bond angles (°) | 0.6 | 0.6 |

A single crystal was used for each structure determination. Values in parentheses are for highest-resolution shell.

**Fig. 1:**
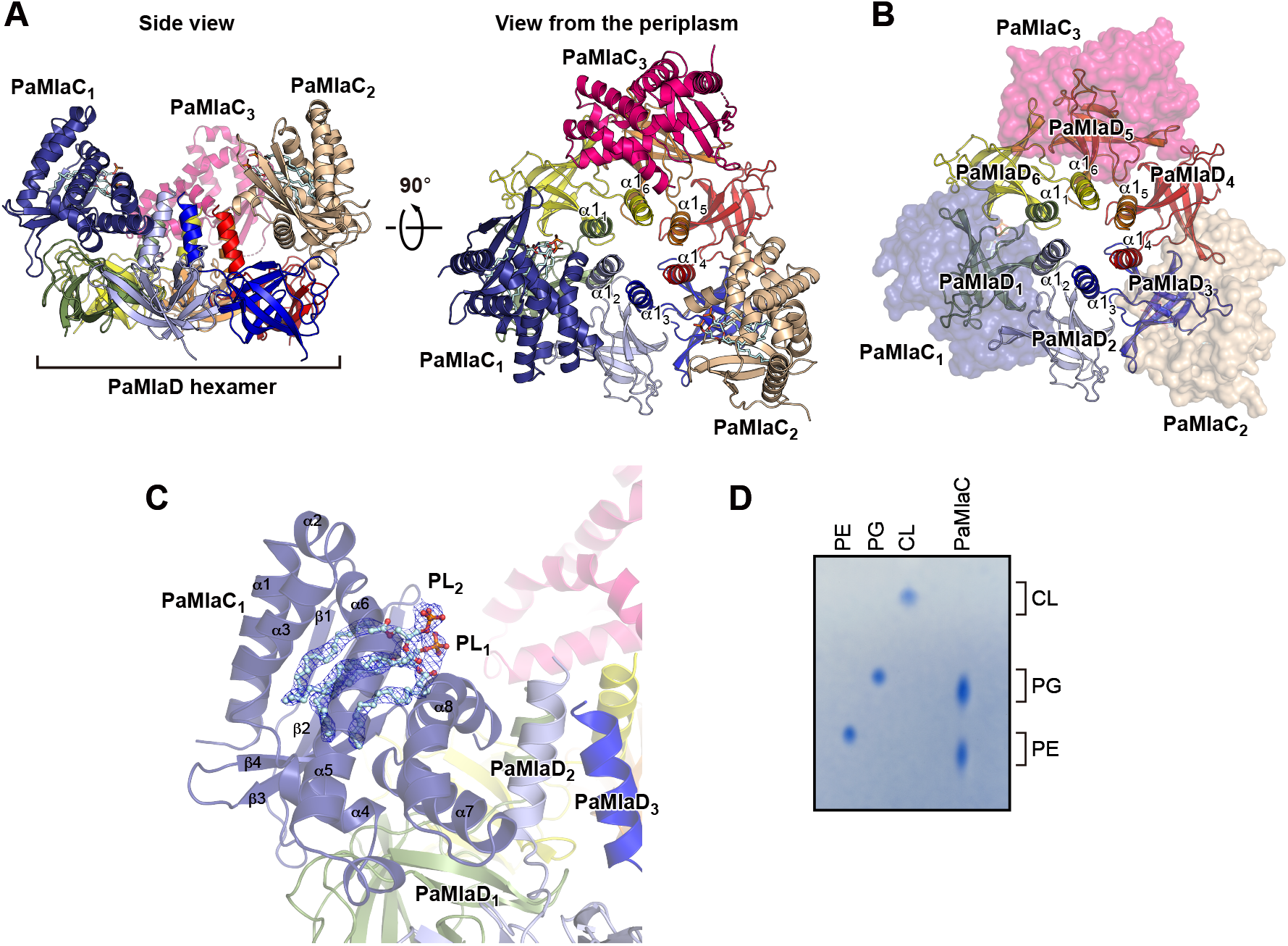
Structure of the PaMlaC–PaMlaD^ΔN36^ complex. (A) Side view (left) and periplasmic view (right) of the ribbon representation of the PaMlaC– PaMlaD^ΔN36^ complex structure. The α1 helices of PaMlaD^ΔN36^ are labeled (α1_1_–α1_6_). PaMlaC_1–3_ molecules are colored in dark blue, wheat, and pink, respectively. PaMlaD_1–6_ molecules are colored in green, light blue, blue, red, orange, and yellow, respectively. (B) Periplasmic view of the semi-transparent surface representation of PaMlaC and the ribbon representation of PaMlaD^ΔN36^. (C) The 2*mF*o – *DF*c electron density map of phospholipid molecules (PL_1_ and PL_2_) bound to PaMlaC_1_, shown as blue mesh and contoured at 1.0 α. (D) Thin-layer chromatography (TLC) analysis of phospholipids extracted from purified PaMlaC and detected by molybdenum blue staining. PE, phosphatidylethanolamine; PG, phosphatidylglycerol; CL, cardiolipin.

### Structural basis of the PaMlaC–PaMlaD interaction

In the structure of the PaMlaC–PaMlaD^ΔN36^ complex, PaMlaC_1_ interacted with PaMlaD_1_ and PaMlaD_2_, burying interface areas of 559 Å^2^ and 409 Å^2^, respectively (calculated using the PISA server^31^), suggesting a more extensive interaction with PaMlaD_1._ The side chain of Arg161 in ^Δ^4 of PaMlaC_1_ forms hydrogen bonds with Asp96 and Asp119 of PaMlaD_1_. In addition, Arg175 in α7 of PaMlaC_1_ forms a hydrogen bond with the main chain of Val116 and is positioned in proximity to Thr98 and Ser115 of PaMlaD_1_ (Fig. 2A and B). Furthermore, α8 of PaMlaC_1_ contacts α1 of PaMlaD_2_, in which Ala209 of PaMlaC_1_ packs against a hydrophobic patch formed by Leu146, Leu147, and Val150 of PaMlaD_2_ (Fig. 2C and D). Structural comparison of the PaMlaD^ΔN36^ hexamer bound to PaMlaC with the MlaD hexamer in the MlaFEDB complex from *P. aeruginosa* (PDB code: 7CH9) showed that the central pore of the PaMlaD hexamer is narrowed due to a 4.6 Å shift of the α1 helix toward the pore upon PaMlaC binding, as observed for PaMlaC_1_ (Supplementary Fig. 3A). These observations suggest that interaction of α8 of PaMlaC with PaMlaD promotes narrowing of the central pore of PaMlaD.

**Fig. 2:**
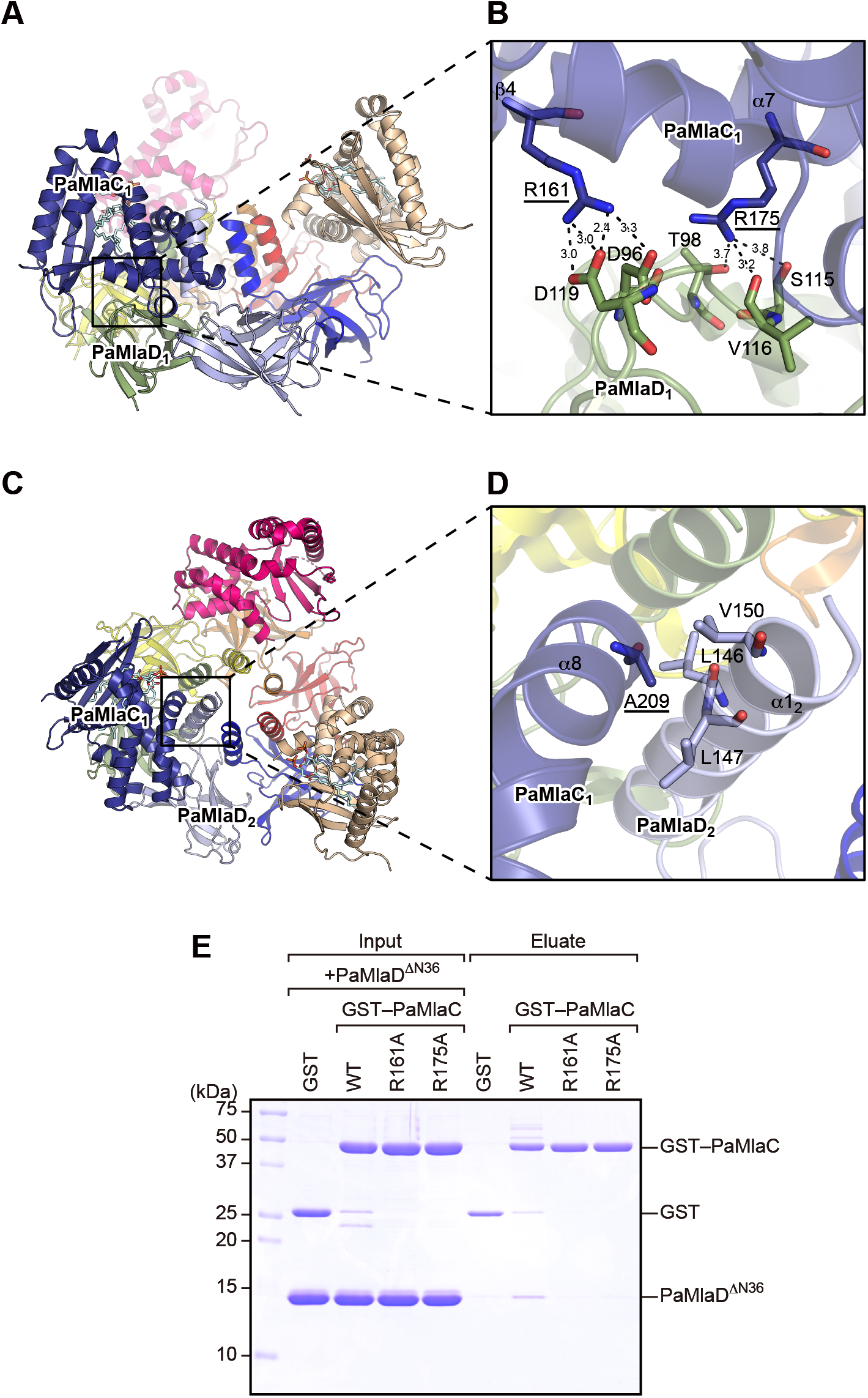
Structural basis of the PaMlaC–PaMlaD^ΔN36^ interaction. (A) The ribbon representation of the PaMlaC–PaMlaD^ΔN36^ complex structure showing the interaction between PaMlaC_1_ and PaMlaD_1_. (B) Magnified view of the interaction surface between PaMlaC_1_ and PaMlaD_1_. Arg161 and Arg175 of PaMlaC_1_ and Asp96, Thr98, Ser115, Val116, and Asp119 of PaMlaD_1_ are shown in stick representation. Residues of PaMlaC_1_ are labeled with underlined text. Dashed lines indicate intermolecular interactions, and the numbers indicate distances (Å). (C) The ribbon representation of the PaMlaC–PaMlaD^ΔN36^ complex structure showing the interaction between PaMlaC_1_ and PaMlaD_2_. (D) Magnified view of the interaction surface between PaMlaC_1_ and PaMlaD_2_. Ala209 of PaMlaC_1_ and Leu146, Leu147, and Val150 of PaMlaD_2_, which are involved in hydrophobic interactions, are shown in stick representation. Ala209 of PaMlaC_1_ is labeled with underlined text. (E) GST pull-down assay showing the interaction between GST–PaMlaC variants and PaMlaD^ΔN36^. GST–PaMlaC variants bound to GST-Accept resin were incubated with PaMlaD^ΔN36^. Proteins bound to the resin were eluted with reduced glutathione and analyzed by SDS-PAGE followed by CBB staining.

To investigate the contribution of Arg161 and Arg175 of PaMlaC to the interaction with PaMlaD, we substituted these two residues with alanine (R161A and R175A), purified N-terminal glutathione S-transferase (GST)-tagged PaMlaC variants, and subjected to an in vitro GST pull-down assay (Fig. 2E). This assay showed that neither the R161A nor R175A variant exhibited detectable association with PaMlaD, indicating that Arg161 and Arg175 of PaMlaC play a critical role in the PaMlaC–PaMlaD interaction.

### Crystal structure of the PaMlaC–PaMlaD^ΔN30^ complex

We next determined the crystal structure of the PaMlaC–PaMlaD^ΔN30^ complex and refined it to 3.46 Å resolution (Table 1). Similarly to the PaMlaC–PaMlaD^ΔN36^ complex, three PaMlaC molecules (PaMlaC_1–3_) were bound to the PaMlaD^ΔN30^ hexamer in a threefold symmetric arrangement in the asymmetric unit (Fig. 3A). No clear phospholipid density was observed within the lipid binding cavities of the PaMlaC molecules, possibly due to the limited resolution. Although the overall structure of the PaMlaC–PaMlaD^ΔN30^ complex was highly similar to that of the PaMlaC–PaMlaD^ΔN36^ complex with an r.m.s.d. of 0.86 Å over 833 Cα atoms, the C-terminal α8 helix of PaMlaC exhibited partial disorder in the PaMlaC–PaMlaD^ΔN30^ complex. Clear electron density for the α8 helix was observed in PaMlaC_1_ and PaMlaC_3_, which adopted similar conformations, whereas the corresponding region (residues 198–215) in PaMlaC_2_ was not well defined, suggesting that this region is flexible and likely adopts multiple conformations. Notably, PaMlaC_2_ was positioned closer to the PaMlaD hexamer than PaMlaC_1_ and PaMlaC_3_, as reflected by a shorter distance between the Cα atoms of Leu173 in PaMlaC and Lys144 in the α1 helix of the nearest PaMlaD subunit to each PaMlaC molecule (6.6 Å for PaMlaC_2_ versus 12.1–12.8 Å for PaMlaC_1_ and PaMlaC_3_) (Fig. 3B, C and Supplementary Fig. 4). Consistent with this, PISA analysis showed that PaMlaC_2_ formed a larger interface with its principal interacting PaMlaD subunit (548.6 Å^2^) than did PaMlaC_1_ and PaMlaC_3_ (402.3 and 312.2 Å^2^, respectively), indicating a more extensive interaction with the PaMlaD hexamer. This closer positioning is likely caused by disordering of the C-terminal α8 helix of PaMlaC. Partial disordering of the α8 helix disrupts its interaction with the α1 helix of PaMlaD, thereby relieving steric hindrance and permitting tighter association with PaMlaD. Consistent with the loss of this interaction, the α1 helix of PaMlaD subunit proximal to PaMlaC_2_ did not adopt a pore-narrowing conformation in this structure (Supplementary Fig. 3B). Importantly, this conformational variability was accompanied by increased exposure of the lipid-binding cavity of PaMlaC (Fig. 3D, E and Supplementary Fig. 4). In the ordered state, the C-terminal α8 helix restricts access to the lipid-binding cavity. These observations suggest that rearrangement of the C-terminal α8 helix is coupled to changes in both the orientation of PaMlaC and the accessibility of its lipid-binding cavity.

**Fig. 3:**
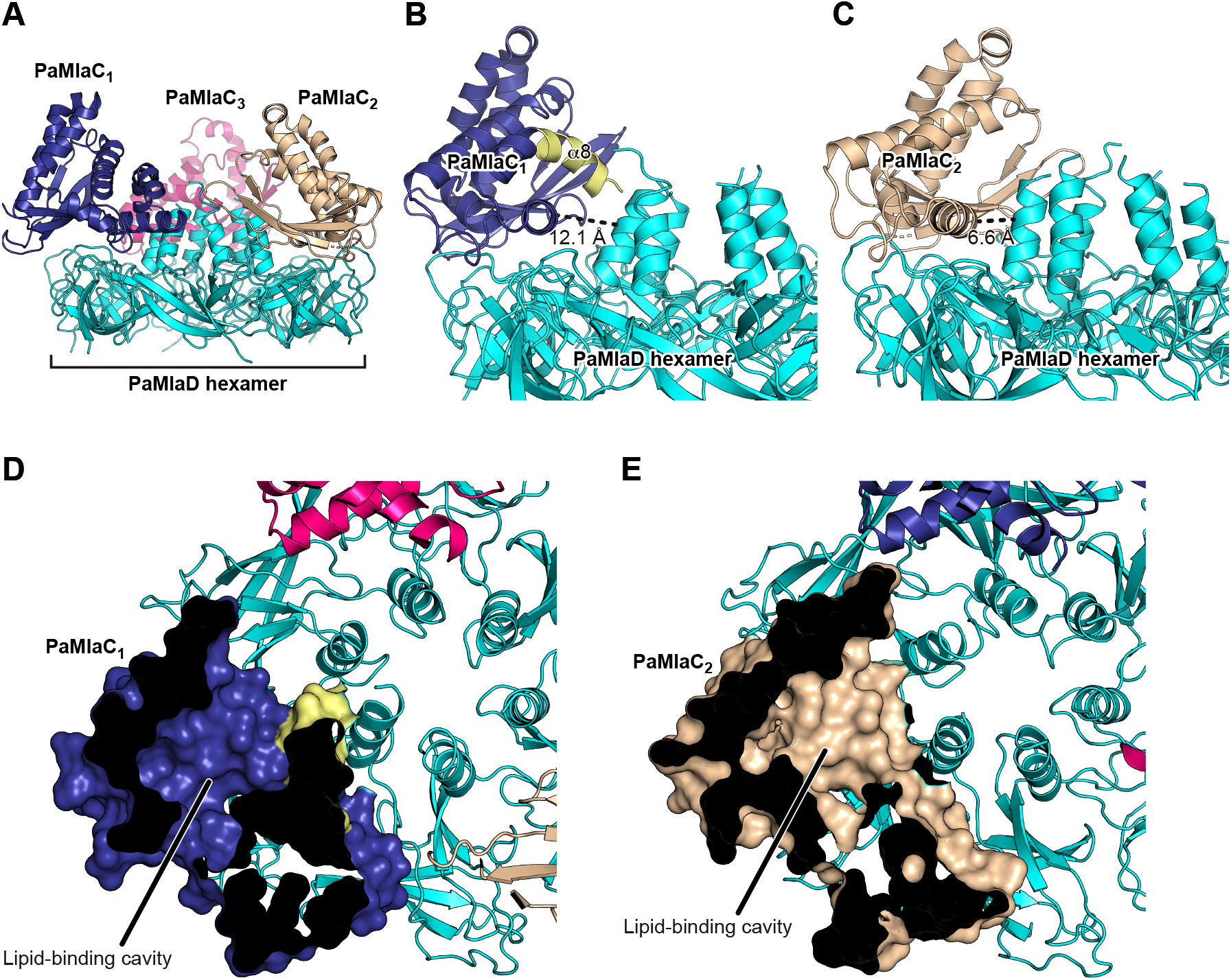
Structure of the PaMlaC–PaMlaD^ΔN30^ complex. (A) Ribbon representation of the PaMlaC–PaMlaD^ΔN30^ complex. PaMlaC_1–3_ molecules are colored in dark blue, wheat, and pink, respectively. The PaMlaD hexamer is colored in cyan. (B) Interaction of PaMlaC_1_ with the PaMlaD hexamer. Part of the C-terminal α8 helix of PaMlaC_1_ is colored in yellow. The distance between the Cα atoms of Leu173 in PaMlaC_1_ and Lys144 in the α1 helix of the nearest PaMlaD subunit is indicated by a dashed line. (C) Interaction of PaMlaC_2_ with the PaMlaD hexamer. The distance between the Cα atoms of Leu173 in PaMlaC_2_ and Lys144 in the α1 helix of the nearest PaMlaD subunit is indicated by a dashed line. (D, E) Cutaway surface representations of PaMlaC_1_ (D) and PaMlaC_2_ (E), showing the lipid-binding cavity.

### The C-terminal region of PaMlaC negatively regulates its interaction with PaMlaD

To examine whether the C-terminal region of PaMlaC influences its interaction with PaMlaD, we generated a GST-tagged PaMlaC variant lacking the C-terminal region (residues 23–197; PaMlaC^ΔC^) and performed in vitro pull-down assays (Fig. 4A). Deletion of the C-terminal region of PaMlaC resulted in increased binding to PaMlaD^ΔN36^ compared to the wild-type protein, indicating that the C-terminal region suppresses the interaction with PaMlaD. To examine whether this regulatory effect is conserved across species, we also performed pull-down assays using *E. coli* MlaC lacking the N-terminal signal peptide (residues 22–211; EcMlaC), an EcMlaC variant lacking the C-terminal region (residues 22–198; EcMlaC^ΔC^), and *E. coli* MlaD lacking the N-terminal signal peptide, the transmembrane helix, and the C-terminal disordered region that is dispensable for function^29^ (residues 37–152; EcMlaD) (Fig. 4B). In contrast to PaMlaC, deletion of the C-terminal region of EcMlaC had a much smaller effect on EcMlaD binding, suggesting that the regulatory role of the C-terminal region differs between species. Consistent with this interpretation, structural comparison revealed that the α8 helix of EcMlaC is shorter than that of PaMlaC (Supplementary Fig. 5A). In addition, in the *E. coli* MlaC–MlaD complex structure, the C-terminal region of MlaC does not interact with the α1 helix of MlaD (Supplementary Fig. 5B). These structural differences suggest that, unlike in *P. aeruginosa*, the C-terminal region of EcMlaC might not significantly contribute to the regulation of its interaction with EcMlaD. To quantitatively evaluate the effect of the C-terminal region on the MlaC–MlaD interaction, we carried out isothermal titration calorimetry (ITC) analysis using purified *P. aeruginosa* proteins. The dissociation constant (*K*_d_) for the interaction between wild-type PaMlaC and PaMlaD^ΔN36^ was determined to be 15.5 μM, whereas truncation of the C-terminal region significantly increased the binding affinity, yielding a *K*_d_ of 5.1 μM (Fig. 4C). The fitted stoichiometry values (*N*) were 0.209 for wild-type PaMlaC and 0.343 for PaMlaC^ΔC^, with no substantial difference between the two variants, suggesting that truncation of the C-terminal region primarily affects binding affinity without substantially altering binding stoichiometry. The fitted stoichiometry values suggest binding of approximately one or at most two PaMlaC molecules per PaMlaD hexamer in solution. These results demonstrate that the C-terminal region of MlaC negatively regulates its interaction with MlaD, at least in *P. aeruginosa*. In contrast, when the same analysis was performed using the *E. coli* proteins, ITC measurements of the EcMlaC–EcMlaD interaction yielded only a very small heat change, preventing reliable determination of the dissociation constant. Taken together with the structural observations in Fig. 3, these findings suggest that conformational changes in the C-terminal α8 helix are likely to influence the relative orientation of PaMlaC and thereby modulate its binding to PaMlaD.

**Fig. 4:**
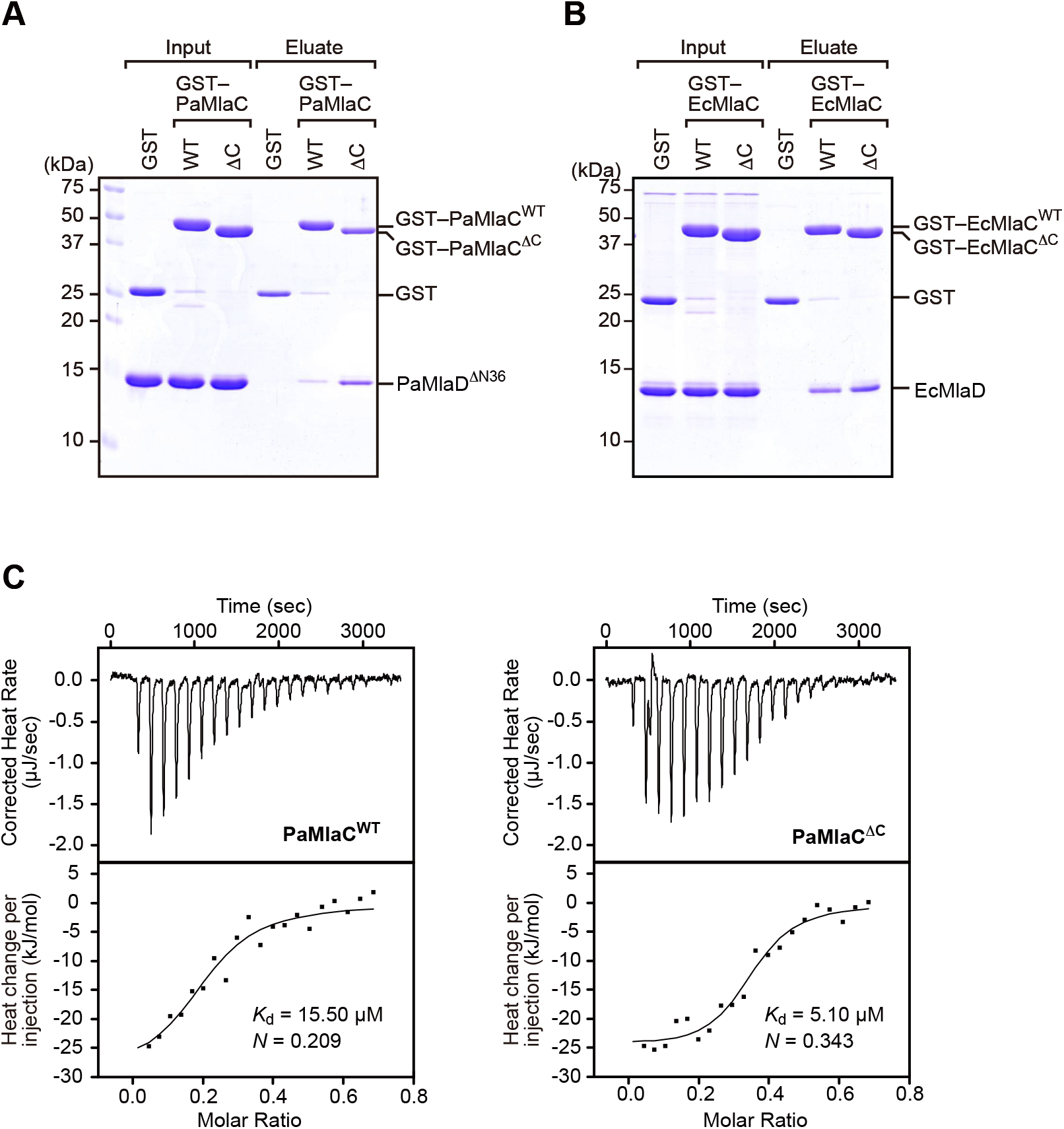
The C-terminal α8 helix of PaMlaC negatively regulates the interaction with PaMlaD. (A) GST pull-down assay showing the interaction between GST–PaMlaC variants and PaMlaD^ΔN36^. GST–PaMlaC^WT^ and GST–PaMlaC^ΔC^ bound to GST-Accept resin were incubated with PaMlaD^ΔN36^. Proteins bound to the resin were eluted with reduced glutathione and analyzed by SDS-PAGE followed by CBB staining. (B) GST pull-down assay showing the interaction between GST–EcMlaC variants and EcMlaD. GST–EcMlaC^WT^ and GST–EcMlaC^ΔC^ bound to GST-Accept resin were incubated with EcMlaD. Proteins bound to the resin were eluted with reduced glutathione and analyzed by SDS-PAGE followed by CBB staining. (C) ITC results obtained by titration of wild-type PaMlaC (left) or PaMlaC^ΔC^ (right) into a solution of PaMlaD^ΔN36^. The dissociation constant (*K*_d_) and binding stoichiometry (*N*) are shown.

### The C-terminal region of MlaC stabilizes phospholipid binding

To investigate whether the C-terminal region of MlaC influences phospholipid binding, we analyzed the phospholipids associated with GST-tagged MlaC proteins following detergent washing. GST-tagged wild-type PaMlaC and PaMlaC^ΔC^ were purified and subjected to washing with n-dodecyl-β-D-maltoside (DDM), and bound phospholipids were extracted and analyzed using TLC followed by ninhydrin staining (Fig. 5A). In the absence of DDM, PE was detected in association with both wild-type PaMlaC and PaMlaC^ΔC^ variants. However, following DDM washing, PE remained associated with the wild-type PaMlaC, whereas it was markedly reduced in PaMlaC^ΔC^. This result indicates that deletion of the C-terminal region decreases the stability of phospholipid binding to PaMlaC. Consistent with this observation, a similar trend was observed for EcMlaC, in which deletion of the C-terminal region also resulted in reduced phospholipid retention (Fig. 5A), suggesting that the role of the C-terminal region in stabilizing phospholipid binding is conserved between *P. aeruginosa* and *E. coli*. To examine whether the reduced phospholipid retention affects phospholipid transport activity, we performed an in vitro phospholipid transport assay using the *E. coli* Mla system, as previously described^13^. Donor liposomes containing NBD-labeled PE (NBD-PE) and Rhodamine-labeled PE (Rhod-PE), reconstituted with the *E. coli* MlaA–OmpC complex, were incubated with wild-type EcMlaC or EcMlaC^ΔC^ and acceptor liposomes reconstituted with the *E. coli* MlaFEDB complex and lacking fluorescent lipids. When NBD-PE and Rhod-PE are present in the same liposome, NBD fluorescence is quenched by Rhodamine, and transfer of NBD-PE to acceptor liposomes results in fluorescence dequenching. In this assay, deletion of the C-terminal region had little effect on overall transport activity compared to the wild-type protein (Supplementary Fig. 6). Although this analysis was limited to the *E. coli* system, the result suggests that the C-terminal region is not strictly required for phospholipid transfer itself. Phospholipid transport activity in the *P. aeruginosa* Mla system could not be evaluated, as MlaA from *P. aeruginosa* was not successfully prepared. Taken together, these findings indicate that the C-terminal region of MlaC contributes to the stable retention of phospholipids within its binding cavity, rather than being strictly required for phospholipid transport.

**Fig. 5:**
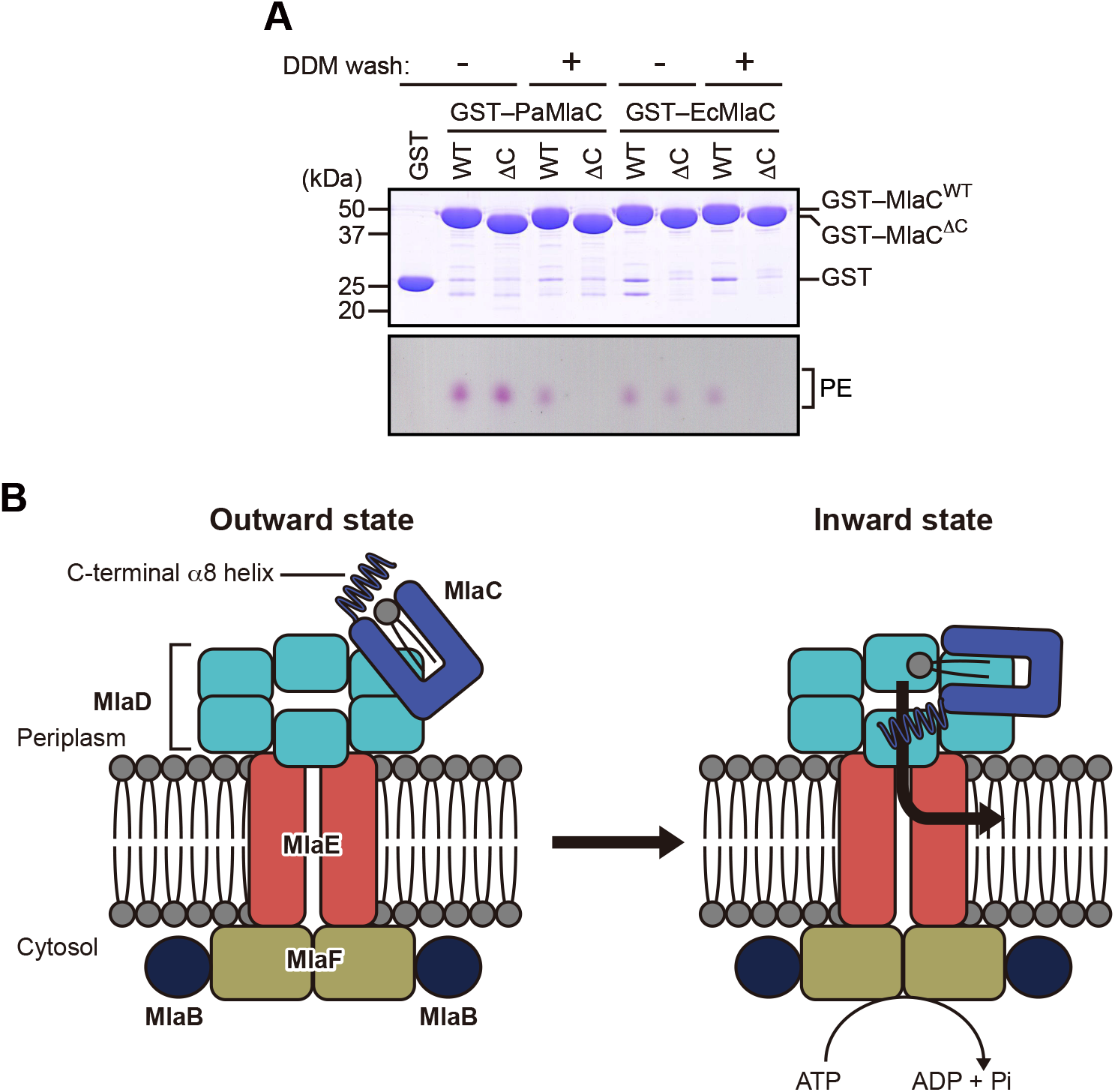
The C-terminal α8 helix of PaMlaC regulates phospholipid binding. (A) Analysis of phospholipids associated with GST-tagged PaMlaC and EcMlaC variants after detergent washing. Purified GST–PaMlaC^WT^, GST–PaMlaC^ΔC^, GST–EcMlaC^WT^, and GST– EcMlaC^ΔC^ immobilized on GST-Accept resin were washed with buffer containing DDM and subsequently eluted with reduced glutathione. Phospholipids associated with the eluted proteins were extracted and analyzed by TLC followed by ninhydrin staining. Protein samples were analyzed by SDS-PAGE followed by CBB staining. (B) Proposed model for conformational changes of MlaC during phospholipid transfer in the *P. aeruginosa* Mla system. In the outward state (left), the C-terminal α8 helix adopts an ordered conformation, and MlaC interacts with the MlaD hexamer in a more distal orientation that limits accessibility of the lipid-binding cavity to the central pore. In the inward state, partial disordering of the α8 helix allows closer association of MlaC with the MlaD hexamer and increased accessibility of the lipid-binding cavity, which may facilitate phospholipid transfer toward the central pore and subsequent ATP-dependent transport by MlaFEDB complex.

## Discussion

In this study, we provide structural and biochemical insights into the mechanism by which the periplasmic lipid carrier MlaC interacts with the MlaD hexamer and may regulate phospholipid transfer within the *P. aeruginosa* Mla pathway. Our findings identify the C-terminal α8 helix of MlaC as a key regulatory element that modulates both its interaction with MlaD and the stability of phospholipid binding, suggesting a potential role in lipid transfer at the MlaC–MlaD interface.

A central observation of this study is that the C-terminal region of MlaC plays a dual role in regulating its structural and interaction properties. Based on two crystal structures of the PaMlaC–PaMlaD complex (Figs. 1 and 3), our analyses suggest that the α8 helix of PaMlaC adopts distinct conformations, which correlate with differences in the orientation of PaMlaC relative to the PaMlaD hexamer and the accessibility of its lipid-binding cavity. When the α8 helix is well defined, it interacts with the α1 helix of PaMlaD, positioning PaMlaC at a defined distance from the PaMlaD surface and restricting access to the lipid-binding cavity (Figs. 2D and 3D). In contrast, when the α8 helix becomes partially disordered, PaMlaC adopts a closer binding mode with PaMlaD, accompanied by increased exposure of the cavity (Fig. 3C and E). Interestingly, these conformational states appear to be consistent with the “outward” and “inward” MlaC–MlaD configurations proposed for the *E. coli* system, where the outward configuration corresponds to a distal position and the inward configuration to a proximal position relative to the central pore of the MlaD hexamer^29^. In our *P. aeruginosa* structures, the conformation with a well-defined α8 helix likely corresponds to the outward state, whereas the partially disordered α8 helix corresponds to the inward state. These structural observations are consistent with our biochemical data showing that deletion of the C-terminal region enhances PaMlaC–PaMlaD interaction (Fig. 4). Our data further suggest that rearrangement of α8 helix toward the inward state promotes phospholipid release by increasing exposure of the lipid-binding cavity (Fig. 5A). Together, these findings support a model in which the C-terminal α8 helix of PaMlaC functions as a regulatory element that modulates the positioning of PaMlaC relative to PaMlaD and the accessibility of its lipid-binding cavity.

Importantly, our results suggest that conformational changes in PaMlaC are coupled to structural rearrangements in PaMlaD (Supplementary Fig. 3). In the structure where PaMlaC adopts an ordered α8 conformation, the α1 helices of the PaMlaD hexamer shift toward the center, narrowing the central pore. In contrast, this rearrangement is not observed when the α8 helix is disordered. This finding suggests that the structural response of PaMlaD depends on the conformational state of PaMlaC rather than on binding alone. The interaction between the α8 helix of PaMlaC and the α1 helix of PaMlaD appears to be central to this coupling, as disorder in α8 weakens this interaction and may allow PaMlaC to approach the PaMlaD surface more closely. Notably, the α8 helix of *Pseudomonas* MlaC adopts a bent conformation that expands the lipid-binding cavity, whereas homologs such as *E. coli* possess a straighter α8 helix associated with a smaller cavity^8,22,23^. This structural difference suggests that the α1 rearrangement observed in PaMlaD may reflect a species-specific mode of interaction adapted to the distinct conformation of PaMlaC.

It is also important to consider the stoichiometry of the MlaC–MlaD interaction. Recent cryo-EM studies have shown that the *E. coli* MlaC–MlaD complex exhibits a stoichiometry of 1:6 or 2:6^29,30^. In contrast, our crystal structures of the PaMlaC–PaMlaD complex display a 3:6 arrangement in the asymmetric unit. However, our ITC analyses suggest that only one or at most two PaMlaC molecules bind to the PaMlaD hexamer in solution (Fig. 4C). These observations indicate that the 3:6 stoichiometry observed in the crystal likely arises from crystallographic packing rather than representing the physiological binding mode.

Based on these findings, we propose a model in which the C-terminal region of MlaC coordinates lipid transfer by regulating both cavity accessibility and interaction with MlaD in the *P. aeruginosa* Mla system (Fig. 5B). In this model, an ordered α8 helix corresponds to an outward state. In this state, MlaC adopts a more distal binding mode relative to the MlaD hexamer and limits access of the lipid-binding cavity to the central pore of the MlaD hexamer. In contrast, partial disordering of this region corresponds to an inward state. In this state, MlaC associates more closely with MlaD and exhibits increased accessibility of the lipid-binding cavity, thereby facilitating lipid transfer toward the central pore. Reformation of the α8 helix following lipid release may promote dissociation from the MlaD hexamer. While this model provides a framework linking local conformational changes in MlaC to the transport cycle, how these states are regulated within the full MlaFEDB complex, particularly in relation to ATP binding and hydrolysis, remains unclear. In addition, the physiological relevance of the multiple binding modes observed here will require further validation. Future studies integrating structural, biochemical, and in vivo approaches will be essential to fully elucidate the mechanism of phospholipid transport in the Mla pathway.

## Methods

### Construction of expression plasmids

DNA fragments were amplified from the genomes of *P. aeruginosa* PAO1 or *E. coli* DH5α. The DNA fragments encoding PaMlaC (residues 23–215), PaMlaC^ΔC^ (residues 23–197), EcMlaC (residues 22–211), and EcMlaC^ΔC^ (residues 22–198) were cloned into the pGEX-6p-1 vector (GE Healthcare) to construct N-terminal GST-fusion protein expression plasmids. The DNA fragments encoding PaMlaD^ΔN30^ (residues 31–157), PaMlaD^ΔN36^ (residues 37–157), and EcMlaD (residues 37–152) were also cloned into a modified pETDuet-1 vector (Novagen) to construct expression plasmids for proteins with a N-terminal His_6_-tag cleavable by human rhinovirus (HRV) 3C protease. The *mlaFEDCB* and *mlaA* coding regions from *E. coli* were amplified from the genome of *E. coli* DH5α and independently cloned into a modified pET21a vector (Novagen) to generate C-terminally His_10_-tagged MlaB and MlaA, respectively. Mutations for amino-acid substitutions were introduced using PCR-based site-directed mutagenesis. All constructs were verified by DNA sequencing.

### Protein expression and purification

All constructs were expressed in *E. coli* C43 (DE3) cells (Lucigen) cultured in Luria–Bertani (LB) medium at 37 °C. When the culture reached an optical density at 600 nm (OD_600_) of approximately 0.8, protein expression was induced with 0.1 mM isopropyl β -D-thiogalactopyranoside. After growth at 25 °C for 18 h, the cells were harvested. For purification of GST-tagged PaMlaC, PaMlaC^ΔC^, EcMlaC, and EcMlaC^ΔC^ proteins, cell pellets were resuspended in buffer A (20 mM Tris-HCl [pH 8.0] and 150 mM NaCl) supplemented with 5 mM dithiothreitol, disrupted by sonication, and centrifuged at 20,000 *× g* for 40 min to remove insoluble debris. The supernatant was applied to a GST-Accept column (Nacalai Tesque) equilibrated with buffer A. The column was washed with buffer A, and GST-tagged proteins were eluted using buffer B (50 mM Tris-HCl [pH 8.0] and 10 mM reduced glutathione). When necessary, the GST tag was cleaved using HRV 3C protease at 4 °C overnight and removed by reloading the sample onto the GST-Accept column. For purification of N-terminally His_6_-tagged PaMlaD^ΔN30^ and PaMlaD^ΔN36^ proteins, cell pellets were resuspended in buffer C (50 mM Tris-HCl [pH 8.0], 500 mM NaCl, and 20 mM imidazole). Cells were disrupted by sonication, and insoluble debris was removed by centrifugation at 20,000 *× g* for 40 min. The supernatant was applied to a Ni-NTA column (Qiagen) equilibrated with buffer C. The column was washed with buffer C, and His_6_-tagged proteins were eluted using buffer D (50 mM Tris-HCl [pH 8.0], 100 mM NaCl, and 250 mM imidazole). The His_6_ tag was cleaved using HRV 3C protease at 4 °C overnight. The affinity-purified proteins were further purified by size-exclusion chromatography using a Superdex 200 Increase column (GE Healthcare) equilibrated with buffer A.

For purification of the MlaFEDB complex carrying a C-terminally His_10_-tagged MlaB and of C-terminally His_10_-tagged MlaA, cells were resuspended in buffer A. After cell disruption by sonication, cell debris was removed by centrifugation at 10,000 *× g* for 10 min and the membrane fraction was collected by ultracentrifugation at 150,000 *× g* for 90 min. The membrane fraction was solubilized in buffer C supplemented with 1.0% dodecyl-β -D-maltopyranoside (DDM) at 4 °C for 60 min. Insoluble components were removed by ultracentrifugation at 150,000 *× g* for 30 min and the supernatant was applied to a Ni-NTA column equilibrated with buffer C supplemented with 0.03% DDM. The column was washed with buffer C supplemented with 0.03% DDM, and bound proteins were eluted using buffer D supplemented with 0.03% DDM. C-terminally His_10_-tagged MlaA was purified together with endogenous OmpC. The MlaFEDB complex and the MlaA–OmpC complex were further purified by size-exclusion chromatography using a Superdex 200 Increase column equilibrated with buffer A supplemented with 0.03% DDM.

### Crystallization and X-ray crystallography

Crystallization trials were performed at 20 °C using the sitting drop vapor diffusion method. For crystallization of the PaMlaC–PaMlaD^ΔN36^ complex, 0.2 μL drops of approximately 15 mg/mL PaMlaC and 9 mg/mL PaMlaD^ΔN36^ in 20 mM Tris-HCl (pH 8.0) and 150 mM NaCl were mixed with an equal amount of reservoir solution consisting of 0.1 M sodium citrate (pH 5.0) and 29% Jeffamine ED-2001 (pH 7.0), and equilibrated against 70 μL of the same reservoir solution through vapor diffusion. For crystallization of the PaMlaC–PaMlaD^ΔN30^ complex, 0.2 μL drops of approximately 15 mg/mL PaMlaC and 9 mg/mL PaMlaD^ΔN30^ in 20 mM Tris-HCl (pH 8.0) and 150 mM NaCl were mixed with an equal amount of reservoir solution consisting of 0.2 M L-proline, 0.1 M HEPES (pH 7.5), and 24% *w/v* polyethylene glycol 1500, and equilibrated against 70 μL of the same reservoir solution through vapor diffusion. Crystals of the PaMlaC–PaMlaD^ΔN36^ complex were flash-cooled in liquid nitrogen without additional cryoprotectant, whereas crystals of the PaMlaC–PaMlaD^ΔN30^ complex were cryoprotected with the reservoir solution supplemented with 20% ethylene glycol prior to flash-cooling in liquid nitrogen. Crystals were maintained in a stream of nitrogen gas at 100 K during X-ray diffraction data collection. Diffraction data were collected at the SPring-8 beamline BL32XU with a 10 *×* 15 μm (width *×* height) microbeam using the helical data collection method. Data collection was performed using the ZOO automated data collection system^32^. The data were processed using the KAMO^33^ and XDS^34^ software. Structures were determined by molecular replacement with PHASER^35^, using the structures of *P. aeruginosa* MlaC (PDB code: 6HSY) and the MlaD hexamer from the *P. aeruginosa* MlaFEDB complex (PDB code: 7CH9) as search models. Further model building was performed manually using COOT^36^, and crystallographic refinement was performed using PHENIX^37^. MolProbity^38^ was used to assess the quality and geometry of structural models. Detailed data collection and processing statistics are shown in Table 1.

### In vitro pull-down assay

Purified GST-tagged PaMlaC or EcMlaC variants (20 μM) and PaMlaD or EcMlaD variants (120 μM) in a total volume of 200 μL were incubated with 50 μL of GST-Accept resin in buffer containing 20 mM Tris-HCl (pH 8.0) and 150 mM NaCl at 4 °C for 60 min. The resin was washed three times with 500 μL of the same buffer, and bound proteins were eluted with 150 μL of 10 mM reduced glutathione in 50 mM Tris-HCl (pH 8.0). Eluted proteins were analyzed by SDS-PAGE followed by Coomassie brilliant blue (CBB) staining.

### TLC analysis

To extract phospholipids bound to PaMlaC, 50 μL of purified PaMlaC (1 mM) in buffer containing 20 mM Tris-HCl (pH 8.0) and 150 mM NaCl was mixed with 750 μL of chloroform/methanol (2:1, *v/v*) and vortexed for 10 min. Subsequently, 100 μL of water was added, and the samples were vortexed again for 10 min. The organic phase was separated by centrifugation at 1,000 × *g* for 2 min, collected, and dried under a stream of N_2_ gas. The resulting lipid film was dissolved in 30 μL of chloroform. Fifteen microliters of the dissolved lipid sample was spotted onto a TLC Silica gel 60 plate (Merck Millipore) and analyzed by TLC using chloroform/methanol/acetic acid (65:25:10, *v/v/v*). Phospholipids were stained with Molybdenum Blue spray reagent (Sigma).

To analyze phospholipids bound to GST-tagged PaMlaC or EcMlaC variants after detergent washing, 500 μL of purified GST-tagged PaMlaC or EcMlaC variants (100 μM) was immobilized onto 100 μL of GST-Accept resin in buffer containing 20 mM Tris-HCl (pH 8.0) and 150 mM NaCl. The resin was washed with 1 mL of the same buffer supplemented with 1% DDM at 4 °C for 60 min. The resin was subsequently washed three times with 500 μL of the same buffer without DDM, and bound proteins were eluted with 200 μL of 10 mM reduced glutathione in 50 mM Tris-HCl (pH 8.0). Phospholipids were extracted from 50 μL of the eluted protein solution as described above. Twenty microliters of the dissolved lipid sample was spotted onto a TLC Silica gel 60 plate (Merck Millipore) and analyzed by TLC using chloroform/methanol/water (65:25:4, v/v/v). Lipid spots were visualized by ninhydrin staining using Ninhydrin-ethanol TS Spray (FUJIFILM Wako).

### Isothermal titration calorimetry (ITC)

All measurements were carried out at 25 °C on a Nano ITC LV calorimeter (TA Instruments), with a stirring rate of 250 rpm. PaMlaC, PaMlaC^ΔC^, and PaMlaD^ΔN36^ were equilibrated in the same buffer (20 mM Tris-HCl pH 8.0 and 150 mM NaCl) by size-exclusion chromatography prior to ITC measurements. PaMlaC or PaMlaC^ΔC^ (1 mM) was titrated into the sample cell containing 170 μL of PaMlaD^ΔN36^ (0.4 mM). The titration consisted of 21 injections of 2 μL each, except for the first injection of 1 μL, with 150 s intervals between injections. To account for the heat of dilution, control titrations of PaMlaC or PaMlaC^ΔC^ into buffer were performed. Data were analyzed using NanoAnalyze software (TA Instruments). Integrated heat data were fitted to a single-site binding model to determine the dissociation constant (*K*_d_) and binding stoichiometry (*N*).

### Liposomes

1,2-dioleoyl-*sn*-glycero-3-phosphoethanolamine (DOPE, 850725C), 1,2-dioleoyl-*sn*-glycero-3-phospho-(1′-rac-glycerol) (DOPG, 840475C), 1′,3′-bis[1,2-dioleoyl-*sn*-glycero-3-phospho-]-*sn*-glycerol (18:1 CL, 710355C), 1-oleoyl-2-(12-[(7-nitro-2-1,3-benzoxadiazol-4-yl)amino]dodecanoyl)-*sn*-glycero-3-phosphoethanolamine (18:1-12:0 NBD-PE, 810146C), 1,2-dioleoyl-*sn*-glycero-3-phosphoethanolamine-*N*-(lissamine rhodamine B sulfonyl) (18:1 Liss-Rhod-PE, 810150C) were obtained from Avanti Polar Lipids. Lipids in stock solutions in chloroform were mixed at the desired molar ratio, and the solvent was evaporated. The lipid film was hydrated in appropriate buffer. The lipid suspension was incubated at room temperature for 30 min and extruded 20 times through a polycarbonate 0.4 μm filter using a mini-extruder (Avanti Polar Lipids).

### Lipid transfer assay

Donor proteoliposomes reconstituted with *E. coli* MlaA–OmpC complex and acceptor proteoliposomes reconstituted with *E. coli* MlaFEDB complex were prepared as previously described^13,30^, with several modifications. For preparation of donor proteoliposomes reconstituted with *E. coli* MlaA–OmpC complex, 700 μL of 0.5 mM liposomes (DOPE/DOPG/18:1 CL/18:1 Liss-Rhod-PE/18:1-12:0 NBD-PE = 60:20:10:2:8) in 20 mM Tris-HCl pH 8.0 and 150 mM NaCl were destabilized by addition of 0.15% DDM and incubated with 100 μL of 10 μM *E. coli* MlaA– OmpC complex at 4 °C for 60 min. For preparation of acceptor proteoliposomes reconstituted with *E. coli* MlaFEDB complex, 700 μL of 2 mM liposomes (DOPE/DOPG/18:1 CL = 70:20:10) in 20 mM Tris-HCl pH 8.0, 150 mM NaCl, 3 mM ATP, and 5 mM MgCl_2_ were destabilized by addition of 0.15% DDM and incubated with 100 μL of 10 μM *E. coli* MlaFEDB complex at 4 °C for 60 min. In both cases, detergent removal and proteoliposome reconstitution were achieved by incubation with 300 mg of SM2 Bio-Beads (Bio-Rad) at 4 °C for 60 min, followed by a second incubation with fresh Bio-Beads under the same conditions.

Lipid transfer assays were carried out by mixing 50 μL each of donor and acceptor proteoliposomes with 50 μL of 100 μM EcMlaC variants in 1850 μL of buffer containing 20 mM Tris-HCl pH 8.0 and 150 mM NaCl. The reaction mixtures were incubated at 30 °C for 2000 s, and NBD fluorescence was monitored using an FP-8500 spectrofluorometer (Jasco) with excitation and emission wavelengths set at 461 and 534 nm, respectively.

## Data availability

The atomic coordinates and structure factors have been deposited in the Protein Data Bank under accession codes 26TP (PaMlaC–PaMlaD^ΔN36^) and 26TQ (PaMlaC–PaMlaD^ΔN30^).

## Acknowledgements

We thank Dr. Yasushi Tamura and Dr. Hisako Kubota-Kawai for their valuable advice, insightful discussions, and support with experimental instruments. This work was supported by the Japan Society for the Promotion of Science KAKENHI (Grant Number JP25K09522 to Y.W.), and Institute for Fermentation, Osaka (to Y.W.). Synchrotron radiation experiments were performed at beamline BL32XU at SPring-8, Japan, with approval from the Japan Synchrotron Radiation Research Institute (Proposal Numbers 2025B2718, 2025A2718, 2024B2757, and 2024A2757).

## Contributions

Conceptualization, Y.W.; Methodology, D.M., S.O., and Y.W.; Investigation, D.M., S.O., and Y.W.; Writing – Original Draft, Y.W.; Writing – Review & Editing, D.M., S.O., and Y.W.; Funding Acquisition, Y.W.; Supervision, Y.W.

