## Supplementary Information for "Structures of the *Pseudomonas aeruginosa* MlaC–MlaD complexes reveal a conformational switch mediated by the C-terminal helix of MlaC"

Supplementary items:

Supplementary Figures 1–6

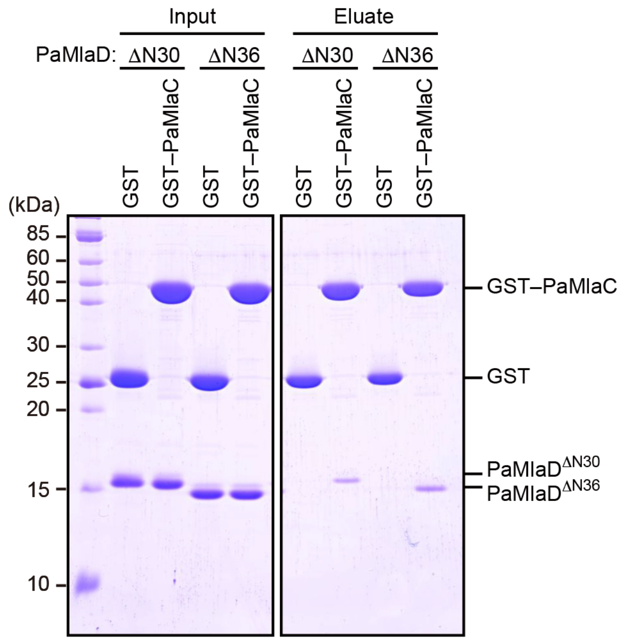


**Supplementary Fig. 1 | Both PaMlaD$^{\Delta N30}$ and PaMlaD$^{\Delta N36}$ interacted with PaMlaC.**

GST pull-down assay showing the interaction between GST–PaMlaC and PaMlaD$^{\Delta N30}$ or PaMlaD$^{\Delta N36}$. GST–PaMlaC bound to GST-Accept resin was incubated with PaMlaD$^{\Delta N30}$ or PaMlaD$^{\Delta N36}$. Proteins bound to the resin were eluted with reduced glutathione and analyzed by SDS-PAGE followed by CBB staining.

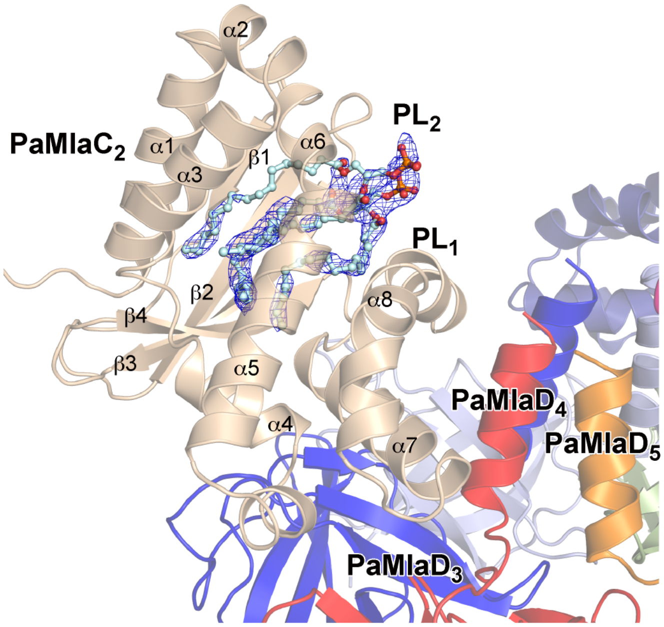


**Supplementary Fig. 2 | Electron density map of phospholipid molecules bound to PaMlaC$_2$ in the PaMlaC–PaMlaD$^{\Delta N36}$ complex.**

The 2*mF*o – *DF*c electron density map of phospholipid molecules (PL$_1$ and PL$_2$) bound to PaMlaC$_2$, shown as blue mesh and contoured at 1.0 σ.

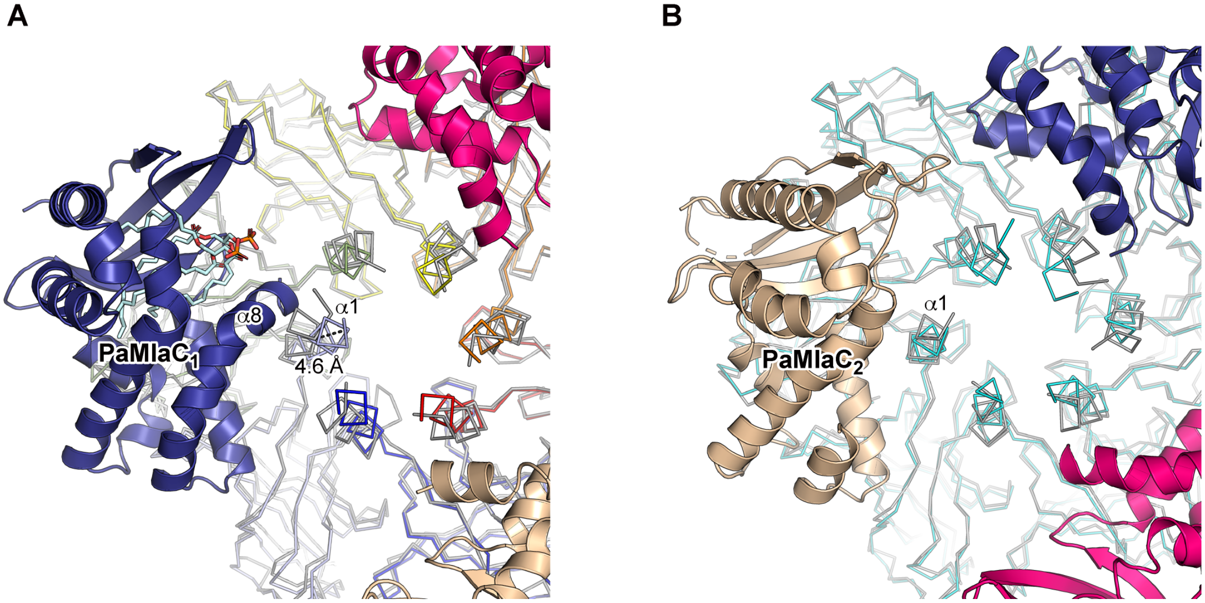


**Supplementary Fig. 3 | Structural comparison of PaMlaD hexamers.**
(A) Superposition of the PaMlaD$^{\Delta N36}$ hexamer in the PaMlaC–PaMlaD$^{\Delta N36}$ complex and the MlaD hexamer in the *P. aeruginosa* MlaFEDB complex (PDB: 7CH9). Upon interaction with the ordered C-terminal α8 helix of PaMlaC$_1$, the α1 helix of PaMlaD$^{\Delta N36}$ adopts a pore-narrowing conformation. The Cα atom of Ser149 is shifted by 4.6 Å relative to the MlaD conformation in the *P. aeruginosa* MlaFEDB structure, as indicated by the dashed line. The color scheme for the PaMlaC–PaMlaD$^{\Delta N36}$ complex is the same as that used in Fig. 1A. (B) Superposition of the PaMlaD$^{\Delta N30}$ hexamer in the PaMlaC–PaMlaD$^{\Delta N30}$ complex and the PaMlaD hexamer in the PaMlaFEDB complex (PDB: 7CH9). In the complex containing a PaMlaC$_2$ molecule with a disordered C-terminal α8 helix, no clear pore-narrowing conformation of the α1 helix of PaMlaD$^{\Delta N30}$ is observed. The color scheme for the PaMlaC–PaMlaD$^{\Delta N30}$ complex is the same as that used in Fig. 3A. In both panels, the MlaD hexamer from the *P. aeruginosa* MlaFEDB structure is colored in gray.

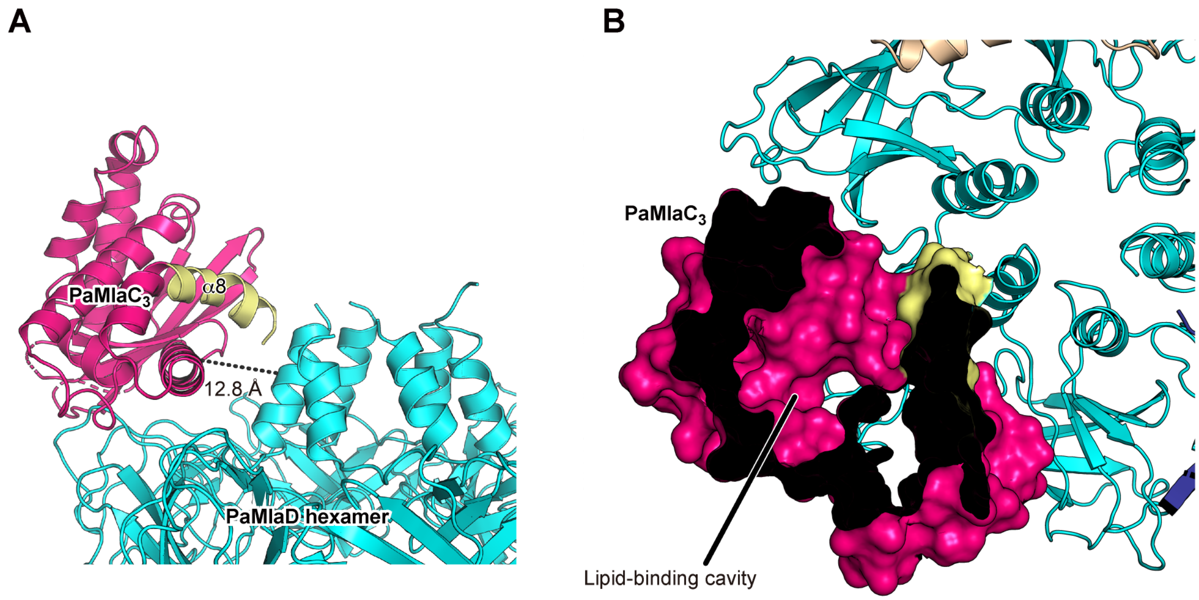


**Supplementary Fig. 4 | Interaction of PaMlaC$_3$ with the PaMlaD hexamer in the PaMlaC–PaMlaD$^{\Delta N30}$ complex.**

(A) Interaction of PaMlaC$_3$ with the PaMlaD hexamer. Part of the C-terminal α8 helix of PaMlaC$_3$ is colored in yellow. The distance between the Cα atoms of Leu173 in PaMlaC$_3$ and Lys144 in the α1 helix of the nearest PaMlaD subunit is indicated by a dashed line. (B) Cutaway surface representations of PaMlaC$_3$, showing the lipid-binding cavity.

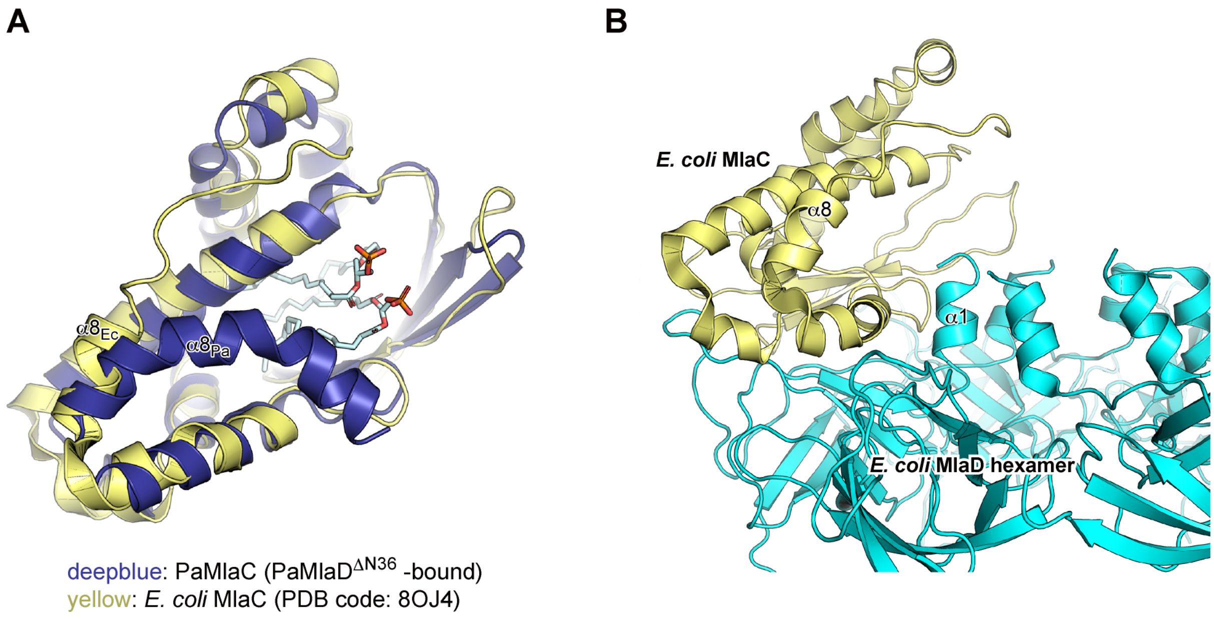


**Supplementary Fig. 5 | Structural comparison between PaMlaC and EcMlaC.**
(A) Superposition of PaMlaC in the PaMlaC–PaMlaD$^{\Delta N36}$ complex and *E. coli* MlaC from the *E. coli* MlaFEDB–MlaC complex (PDB: 8OJ4). PaMlaC and *E. coli* MlaC are colored in dark blue and yellow, respectively. α8 helices of PaMlaC and *E. coli* MlaC are indicated as $\alpha8_{Pa}$ and $\alpha8_{Ec}$, respectively. (B) Interaction of *E. coli* MlaC with the *E. coli* MlaD hexamer in the cryo-EM structure of the *E. coli* MlaFEDB–MlaC complex (PDB: 8OJ4).

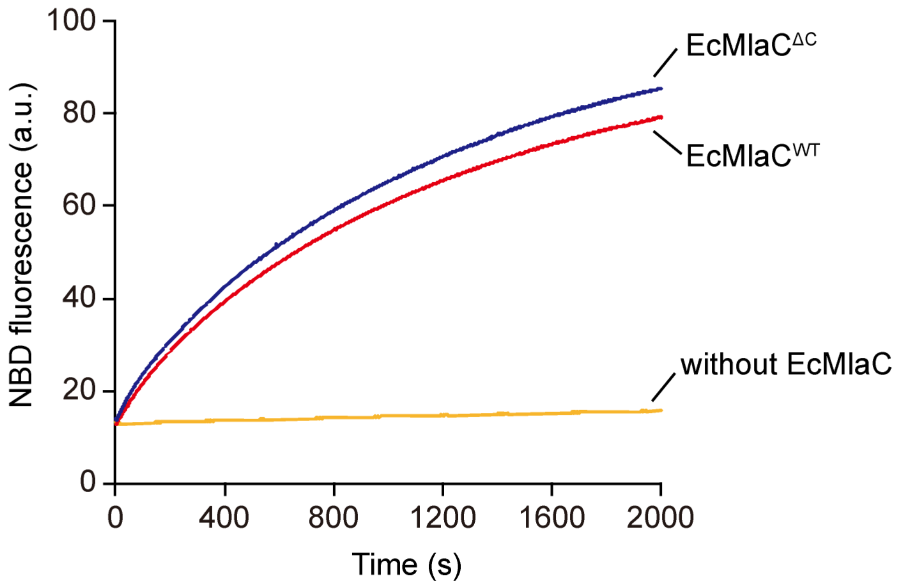


**Supplementary Fig. 6 | Phospholipid transfer assay.**
Phospholipid transfer activity of wild type EcMlaC (EcMlaC$^{WT}$) or EcMlaC$^{\Delta C}$ from MlaA–OmpC proteoliposomes to MlaFEDB proteoliposomes was measured at 30 °C using a fluorescence-based phospholipid transfer assay (see Methods).
